# Proximity proteomics identifies cancer cell membrane *cis*-molecular complex as a potential cancer target

**DOI:** 10.1101/455501

**Authors:** Norihiro Kotani, Arisa Yamaguchi, Tomoko Ohnishi, Ryusuke Kuwahara, Takanari Nakano, Yuka Nakano, Yui Ida, Takayuki Murakoshi, Koichi Honke

**Affiliations:** From the Department of Biochemistry, Saitama Medical University, 38 Morohongo, Moroyama-machi, Iruma-gun, Saitama 350-0495, Japan; From the Department of Biochemistry, Kochi University Medical School, Nankoku, Kochi 783-8505, Japan; From the Quantum Wave Microscopy Group, Okinawa Institute of Science and Technology Graduate University (OIST), Kunigami-gun, Okinawa 904-0495, Japan

**Keywords:** Membrane protein, lipid raft, lung cancer, cancer therapy, protein-protein-interaction.

## Abstract

Cancer-specific antigens expressed in the cell membrane have been used as targets for several molecular targeted strategies in recent years with remarkable success. To develop more effective cancer treatments, novel targets and strategies for targeted therapies are needed. Here, we examined the cancer cell membrane-resident “*cis*-bimolecular complex” as a possible cancer target (*cis*-<u>bi</u>molecular <u>ca</u>ncer <u>t</u>arget: BiCAT) using proximity proteomics, a technique that has attracted attention in recent years. BiCATs were detected using a previously developed method, termed the enzyme-mediated activation of radical source (EMARS), to label the components proximal to a given cell membrane molecule. EMARS analysis identified some BiCATs, such as close homolog of L1 (CHL1), fibroblast growth factor 3 (FGFR3) and α2 integrin, which are commonly expressed in mouse primary lung cancer cells and human lung squamous cell carcinoma cells. Analysis of cancer specimens from 55 lung cancer patients revealed that approximately half of patients were positive for these BiCATs. *In vitro* simulation of effective drug combinations used for multiple drug treatment strategy was performed using reagents targeted to BiCAT molecules. The combination treatment based on BiCAT information moderately suppressed cancer cell proliferation compared with single administration, suggesting that the information about BiCATs in cancer cells is profitable for the appropriate selection of the combination among molecular targeted reagents. Thus, BiCAT has the possibility to make a contribution to several molecular targeted strategies in future.

---

Molecular targeted strategies using specific targets in cancer cells have been widely used in the field of drug discovery (1, 2), drug delivery (3), drug administration (4) and diagnosis (5, 6). The application of these treatment strategies has resulted in good outcomes in terms of cancer diagnosis and treatment and received a certain praises resulting in further competition. However, there are still many difficulties in the development of novel cancer targets that show acceptable efficacy. It is, therefore, necessary to identify novel and potentially effective cancer targets and targeting strategies.

Many molecular targeted strategies have been developed against cell surface (membrane) proteins such as receptor tyrosine kinases (RTKs), which are involved in cell proliferation and differentiation. Previous studies showed that cell surface (membrane) proteins non-randomly forms a hetero-complex accompanied by the fluidity of biological membranes (7). In particular, regions in the membrane with high concentration of specific molecular complexes together with specific lipids are “lipid rafts”. These lipid rafts in the cellular membrane serve as a platform for intracellular signaling and are also involved in various biological phenomena (7). In addition, research in drug discovery and treatment against several diseases has focused on lipid rafts (3, 8, 9). Thus, it is essential to identify the molecules that form *cis*-molecular complexes in the cell membrane, especially cancer cell-specific complexes, with the aim of applying these findings to targeted strategies.

Proximity proteomics (10–13) has recently been used as a method to analyze molecular complexes. We developed a simple and physiological method, called the <u>E</u>nzyme-<u>M</u>ediated <u>A</u>ctivation of <u>R</u>adical Source (EMARS) method (14), which uses horseradish peroxidase (HRP)-induced radicals derived from arylazide or tyramide compounds (15). The EMARS radicals attack and form covalent bonds with the proteins in the proximity of the HRP (*e.g.* radicals from arylazide; approximately 200–300 nm (14), from tyramide; approx. 20 nm (16)) because the generated radicals immediately react with surrounding water molecules and disappear when near HRP. Therefore, the bimolecular partner proteins that interact and assemble with an overexpressed given membrane protein, which was selected based on cDNA microarray data, could be labeled only with arylazide or tyramide compounds under physiological conditions (Fig. S1). The labeled proteins can subsequently be analyzed using an antibody array and/or typical proteome strategy (17). The EMARS method has been applied to various studies on molecular complexes on the cell membrane (18–24).

Here, we propose a “*cis-*bimolecular complex” on the cell membrane as a new type of cancer target (*cis*-bimolecular cancer target, hereinafter referred to as BiCAT) that was identified in pursuit of diversifying molecular targeted strategies. We used the EMARS method in *EML4-ALK* mouse primary lung cancer cells (*EML4-ALK* primary cells) and LK2 human lung squamous cell carcinoma cell line to identify several BiCATs. These BiCATs were also expressed in pathological specimens derived from lung cancer patients.

## Results

### CHL1 is a suitable molecule for BiCAT analysis in EML4-ALK transgenic mouse

The overall scheme of BiCAT analysis for cancer cells is summarized in Fig. 1. The first step is to identify the overexpressed molecules in cancer cell membranes by cDNA array and prepare the EMARS probe. Next, EMARS is performed in (primary) cancer cells and tissues to identify BiCAT partner molecules associated with overexpressed molecules by proteome analysis. BiCAT information is used for further applications (*e.g.* drug design and the simulation of appropriate drug combination for multi-drug administration as described later).

**Fig. 1.**
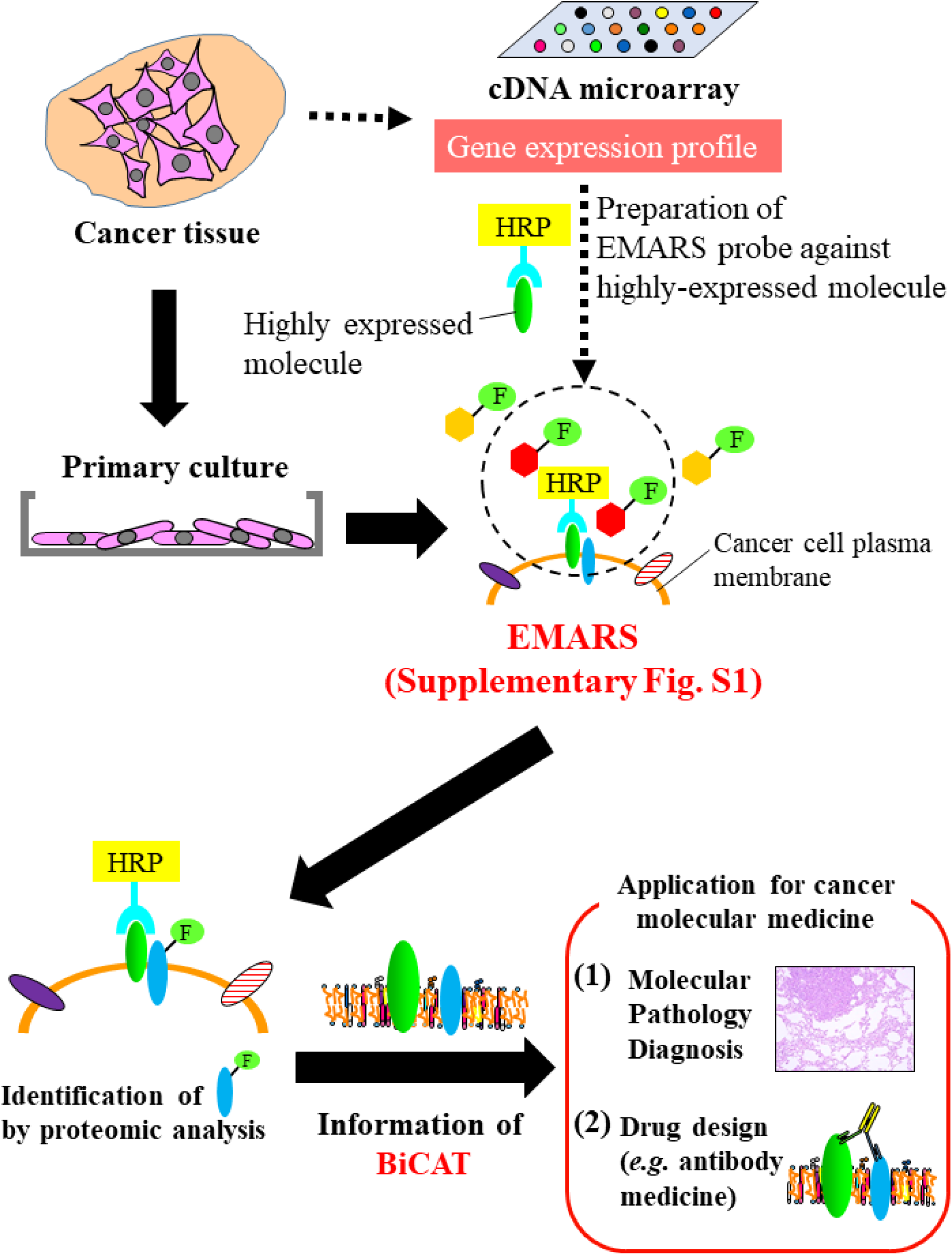
Overview of BiCAT analysis for cancer cell membrane. Schematic illustration of BiCAT analysis. Before the EMARS method, the cancer tissues from *EML4-ALK* transgenic mouse were applied to cDNA microarray analysis, and primary cell inoculation and cultivation.

We used the transgenic mouse of the onco-fusion gene, *Echinoderm microtubule-associated protein-like 4 and Anaplastic lymphoma kinase* (*EML4-ALK*) (25, 26), which causes spontaneously occurring lung cancer with early onset, since it is suitable for biochemical experiments. We first performed gene expression analysis in both lung tumor and normal tissues from *EML4-ALK* transgenic mice (Fig. 2A) by whole mouse cDNA microarray to identify highly expressed membrane molecules in lung tumors (Table S1). We selected four genes (*Gjb4*, *MMP13*, *CHL1* and *Claudin 2*) that were overexpressed in lung tumor tissues as candidate membrane proteins. Reverse transcription PCR revealed that these genes were strongly expressed in lung cancer tumors compared with normal tissue (Fig. 2B) regardless of sex and age. CHL1 expression was detected in tumor slices (Fig. 2C) and in lysates from lung tumors by western blot (Fig. 2D), but not in normal lung tissue. CHL1 was reported as an overexpressed gene in human lung carcinoma tissue (27) and we thus selected CHL1 for subsequent analysis.

**Fig. 2.**
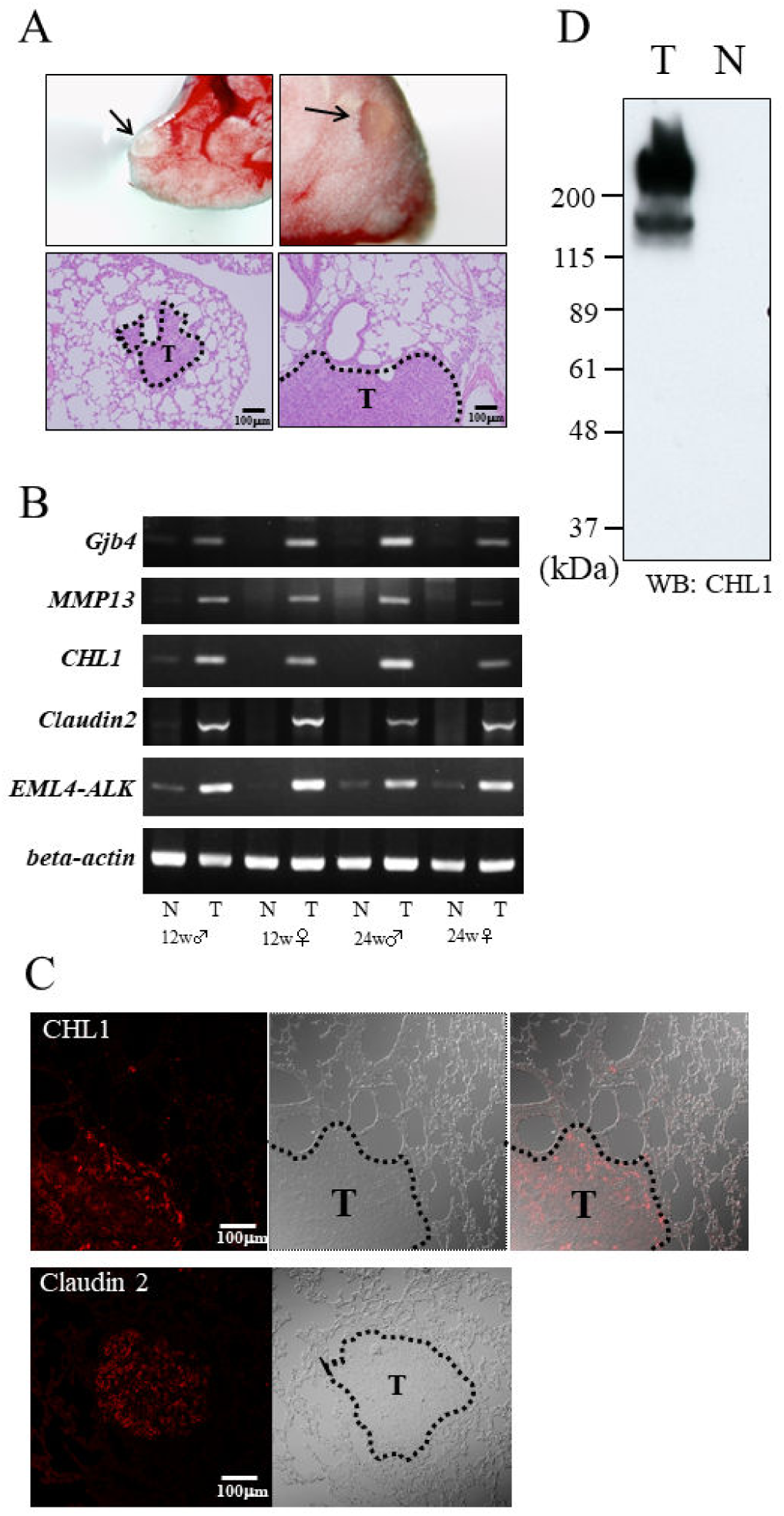
CHL1 expression in lung tumor from *EML4-ALK* transgenic mouse. (*A*) *EML4-ALK* transgenic mouse lung cancers (*Arrows*). Two representative tumor formations in the lung (*Upper panel*) and HE staining of cancer tissue (*Lower panel*; indicated as the dotted area of “T”). Scale bar; 100 μm (*B*) RT-PCR analyses of *Gjb4, MMP13, CHL1, Claudin2,* and *EML4-ALK* mRNAs show potent expression in lung cancer tissue. The mRNA signals of beta-actin were used as a housekeeping gene control. Tissues derived from 12 and 24 weeks old male and female mice were used for the analysis, respectively. N: Normal tissue T: Tumor tissue. (*C*) Immunohistochemical staining of lung tissues from *EML4-ALK* transgenic mouse. CHL1 staining (*Upper panel*) and Claudin2 staining (*Lower panel*) were performed using anti-CHL1 and anti-Claudin2 antibodies with DIC images. Tumor tissue (*T*) was indicated as the dotted area. (*D*) Protein expression of CHL1 in cancer tissue. Tissue lysate from lung cancer tissue and normal tissue were subjected to western blot analysis using mouse CHL1 antibody. N: Normal tissue T: Tumor tissue.

### Partner molecules constituting BiCAT with CHL1

We next used primary cancer cells derived from lung cancer tissue of *EML4-ALK* transgenic mice (Fig. 3A) in EMARS reactions with CHL1 probe (Fig. S2). HRP-conjugated cholera toxin subunit B (CTxB probe), which is the cognitive molecule against ganglioside GM1 as a lipid raft marker (28), was used as a positive control. Using arylazide reagent, the CTxB probe sample generated strong signals; however, moderate signals were observed with the CHL1 probe (Fig. 3B). Weak non-specific signals were observed in the negative control. In contrast, EMARS reaction using tyramide reagent showed clear signals in the CHL1 probe sample, with very faint signals in the negative control (Fig. 3C), suggesting that tyramide-fluorescein reagent was suitable for this study in terms of specificity and sensitivity.

**Fig. 3.**
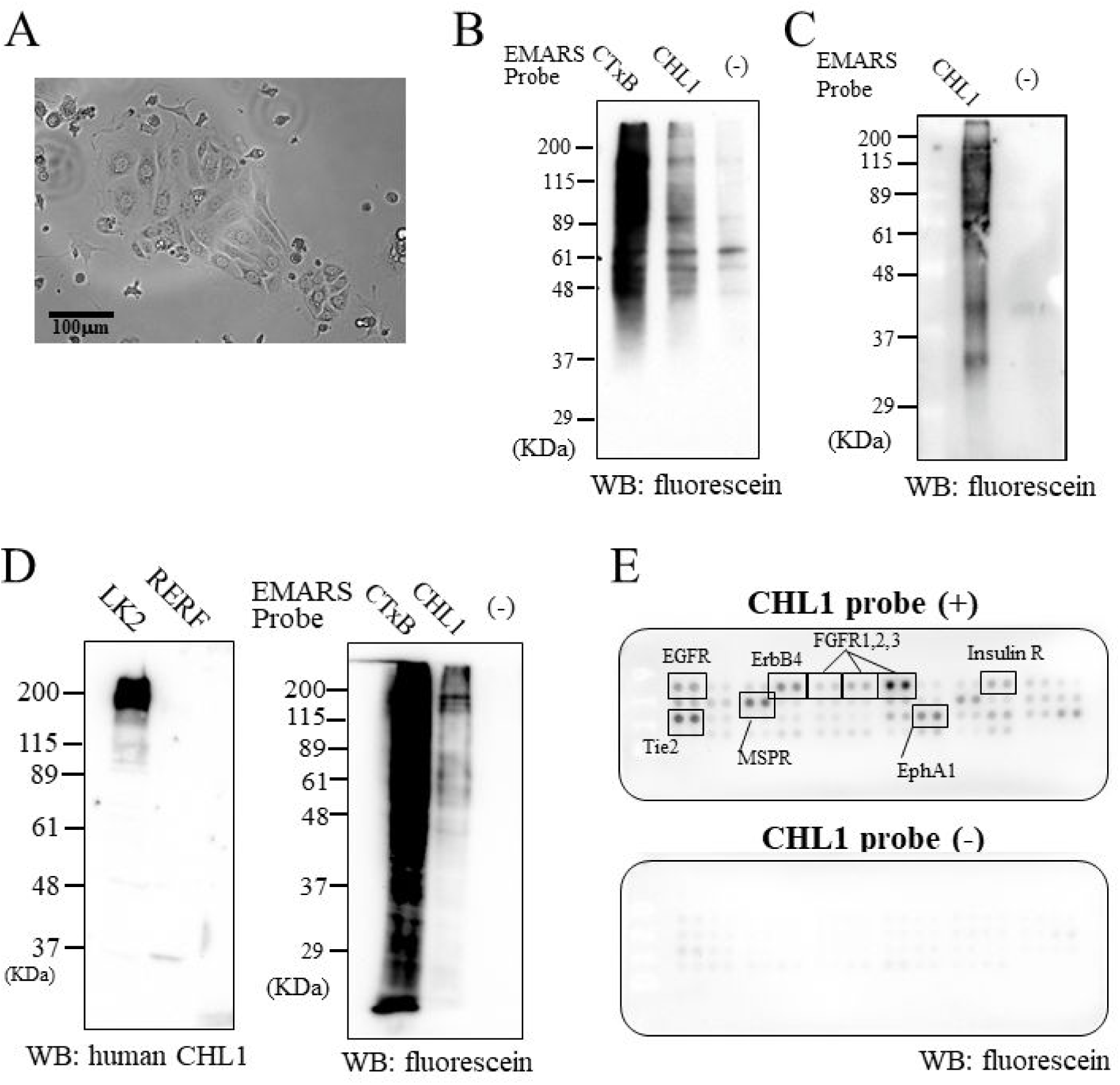
BiCAT analysis for cultured cancer cells. (*A*) Representative image of *EML4-ALK* primary cells. (*B, C*) Partner molecules with CHL1 in *EML4-ALK* primary cells were labeled with fluorescein-arylazide (*B*) and fluorescein-tyramide (*C*) reagent. EMARS products were respectively subjected to western blot analysis followed by the staining using anti-fluorescein antibody. “CTxB” indicates as the positive control sample using CTxB probe, “CHL1” as the samples using CHL1 probe, and “(-)” as negative control samples (no probe). (*D*) EMARS products labeled with fluorescein-tyramide in LK2 cells. Protein expression level of CHL1 in LK2 and RERF cells (*Left column*). EMARS products by CTxB and human CHL1 probes (*Right column*). Abbreviations were same as Fig. 3C. (*E*) Human RTK antibody array analysis of EMARS products from LK2 cells. EMARS samples were applied to Human RTK antibody array according to the manufacture's instruments. “CHL1 probe (+)” indicates as the sample using CHL1 probe, and “CHL1 probe (-)” as negative control samples (no probe). The proteins correspond to positive RTKs were indicated on the array data.

In the human lung carcinoma cell lines, CHL1 protein was expressed in LK2 cells but not in RERF cells (Fig. 3D). EMARS reaction in LK2 cells indicated that both CTxB and human CHL1 probe sample contained fluorescein-labeled proteins indicated as the partner molecules with CHL1 but not in negative control sample (Fig. 3D). The EMARS products were subsequently used for proteomic analysis with mass spectrometry. The identified membrane (-bound) proteins as high possibility candidates for bimolecular partner molecules with CHL1 are summarized in Table S2 (raw data are in Tables S3–S6). The mass spectrometry analysis occupies a large weight for BiCAT analysis, but it is also a useful tool for antibody arrays in terms of its simplicity and sensitivity, especially for low expression molecules in protein lysates. The human RTK antibody array analysis for LK2 cells demonstrated that EMARS products using CHL1 probe contained some RTKs consisting of ErbB and FGFR family members (Fig. 3E). We selected six membrane (-bound) proteins, α2 integrin, β1 integrin, FGFR3, Na/K ATPase, clusterin and contactin1, as bimolecular partners with CHL1. We examined their protein expression in three types of cancer cells and tissue (LK2 cells, *EML4-ALK* primary cells (EML4-PC) and *EML4-ALK* cancer tumor tissue (EML4-T)) by western blot. Although we observed differences in the expression levels of the proteins among the cell types, these proteins, except for contactin1, were endogenously expressed in all three types of cancer cells (Fig. S3A). The multiple bands of clusterin indicated some clusterin isoforms (29). To elucidate whether these proteins labeled with fluorescein, purified fluorescein-labeled EMARS products were subjected to immunopurification and western blot analysis. The fluorescein-labeled α2 integrin, FGFR3 and contactin-1 were detected in the EMARS products from *EML4-ALK* primary cells and LK2 cells (Fig. S3B).

One of the advantages of BiCAT involved in specificity to targeting seemed to be the double assignment using two individual molecules. The best selection for BiCAT molecules are those that are highly expressed in the target sample but not co-expressed in other cells, organs, or tissues. Because they should be followed the basic cancer antigen concept (a large amount of antigen and cancer cells and tissue-specific distribution). We can use the BioGPS database (30), the Human Protein Atlas database (31) and the Human Proteome Map database (32), which provide gene and protein expression profiles classified by organs and tissues for appropriate selection if needed (Fig. S4 and S5). These protein expression profiles demonstrated that the most appropriate CHL1 BiCATs for specific targeting seemed to be CHL1-contactin-1 and CHL1-FGFR3.

### Localization of BiCATs in cancer cell membrane

We next examined whether the identified BiCATs co-localized in the cell membrane. Confocal microscopy showed that α2 integrin (Fig. S6A), β1 integrin (Fig. S6B), clusterin (Fig. S6C), Na/K ATPase (Fig. S6D), FGFR3 (Fig. S6E) and contactin1 (Fig. S6F) co-localized with CHL1, demonstrating that CHL1 and these partner molecules formed BiCATs with each other under an optical microscope. Electron microscopy using LK2 cells (Fig. 4A–C) demonstrated that high levels of gold colloid signals of CHL1 (10 nm particles) and partner molecules (5 nm particles) were located in proximity on the cell membrane. The proteins were located relatively close to each other, with an interval of approximately 10 to 50 nm. Moreover, many 5 nm and 10 nm particles were observed in cellular vesicles (Fig. 4A–C), demonstrating that BiCATs existed not only in cell membranes but also in vesicular membranes.

**Fig. 4.**
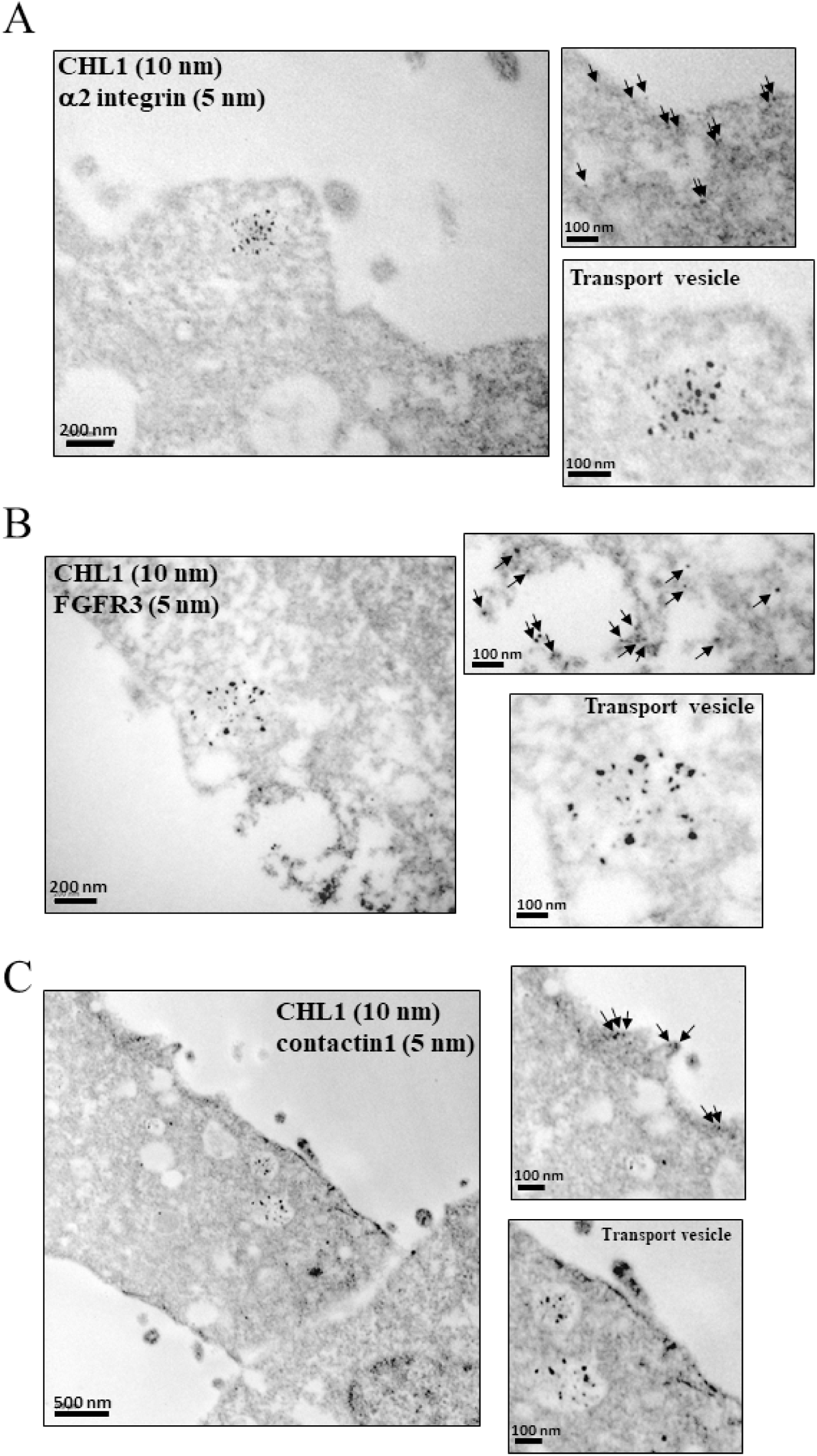
BiCATs located in lung cancer cell membranes and cellular vesicles. (*A to C*) Morphological observation of BiCATs in LK2 cells using electron microscopy. Cultured LK2 cells were fixed and co-stained with CHL1 (indicated as 10 nm particles) and partner molecules identified in cell membrane. α2 integrin (*A*), FGFR3 (*B*), and contactin1 (*C*) were indicated as 5 nm particles. *Arrows* indicate as the locations of gold particles. Scale bar; 100 to 500 nm.

### BiCATs in pathological specimens from lung cancer patients

Histopathological specimens derived from 55 mongoloid cases of lung cancer patients were stained with antibodies against CHL1, α2 integrin, FGFR3 and contactin1. We first performed analysis under low magnification (×5 objective) to detect co-localization signals between CHL1 and partner molecules indicated as BiCATs. CHL1 and the partner molecules were independently expressed in most tissues among 55 cases of lung cancer patients, but did not show the same expression patterns among patients (Fig. S7A–S7C). Both whole and local expressions in the sections were observed. Representative imaging of CHL1, α2 integrin, FGFR3 and contactin1 expression is shown in Fig. 5. Although non-specific signals were observed in all specimens, some tumor specimens had clear or moderate co-localization signals of CHL1-α2 integrin, CHL1-FGFR3 and CHL1-contactin1in the specific areas where cancer cells might be densely packed. The co-localization area of each tumor specimen was then observed under high magnification (×20 objective) and clear co-localization signals as BiCATs were found in specific cancer cells. Next, we classified the tumor specimens into two groups: “positive BiCATs” and “negative BiCATs” as described in *Supporting experimental procedures* and Fig. S7D. Twenty-five cases of CHL1-α2 integrin (45%), 29 cases of CHL1-FGFR3 (53%), and 30 cases of CHL1-contactin1 (55%) were classified as “positive BiCATs” (Fig. S7E and S7F). However, there were no significant relationships between positive staining and patient basic information (age, sex, clinical stage, smoking history; (Fig. S7G–S7I).

**Fig. 5.**
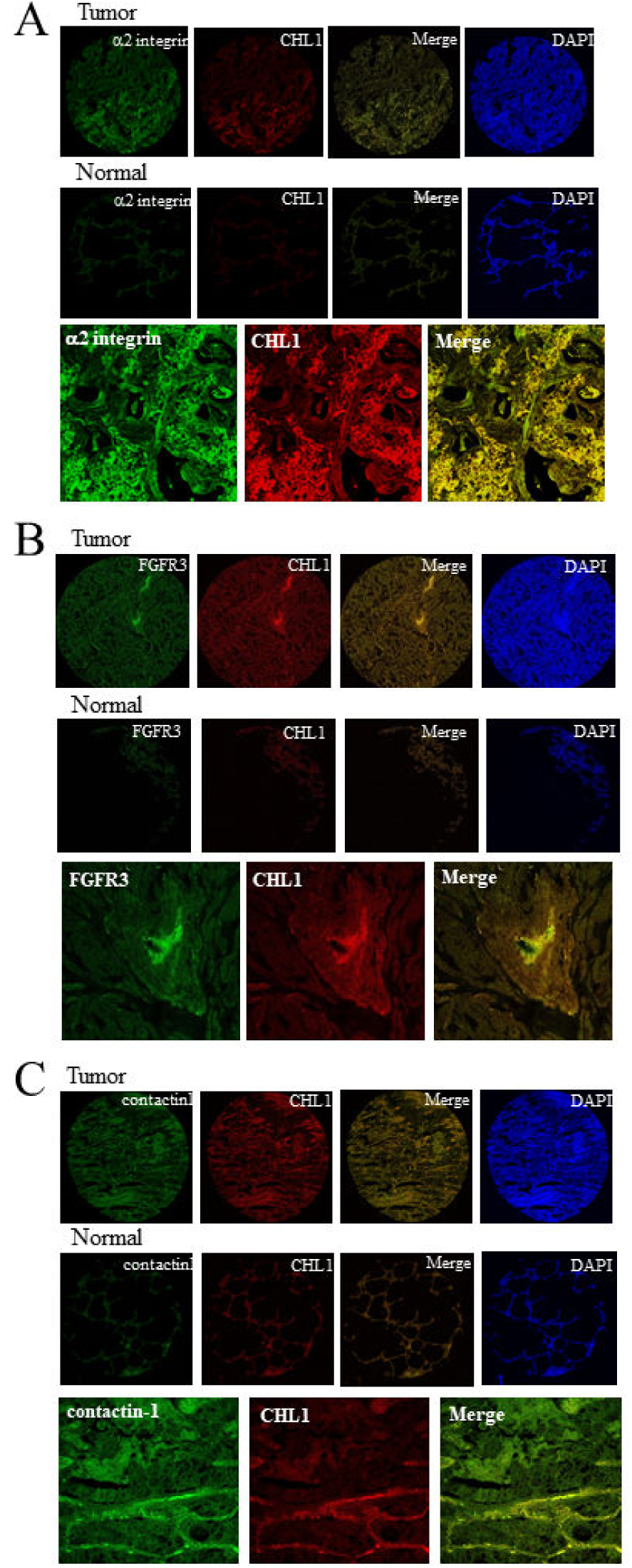
BiCATs located in the pathological specimens from lung cancer patients. (*A to C*) Representative images of BiCAT-positive specimens from 55 cases of lung cancer patients. The lung cancer specimens were co-stained with anti-CHL1 antibody (red) and the antibodies recognizing α2 integrin (*A*), FGFR3 (*B*), contactin1 (*C*) (green), respectively. DAPI solution was used for the nuclear DNA staining. Then, the resulting specimens were observed with confocal microscopy (x5 objective). Both tumor tissues (*Upper panel*) and normal tissue (*Middle panel*) were stained under the same conditions. (*Lower panel*) Representative images at high magnification observation (x20 objective) in part of the positive region of BiCATs indicated as the merged area (yellow).

### In vitro simulation of effective drug combination used for multiple drug treatment strategy based on BiCAT information

Using BiCAT information, we tried a new approach, which was intended for the improvement of multiple drug therapy (33), involving cancer cell proliferation inhibition by multiple antibody/inhibitor administration (anti-CHL1 antibody, FGFR3 inhibitor, and α2β1 integrin inhibitor) against the molecules constituting BiCATs. We compared the efficacy between single and double administration of these antibody/inhibitors under three administration protocols (once daily, and every-other-day protocols; Fig. 6 and Fig. S8) using *in vitro* proliferation inhibition assays. Statistical analyses were performed by both Tukey’s and Dunnett's multiple test (Fig. 6; Dunnett's multiple test). The results of Tukey’s analysis are summarized in Table S7. The efficiency between single and double administration was examined using the statistical significance from the results of Tukey’s analysis.

**Fig. 6.**
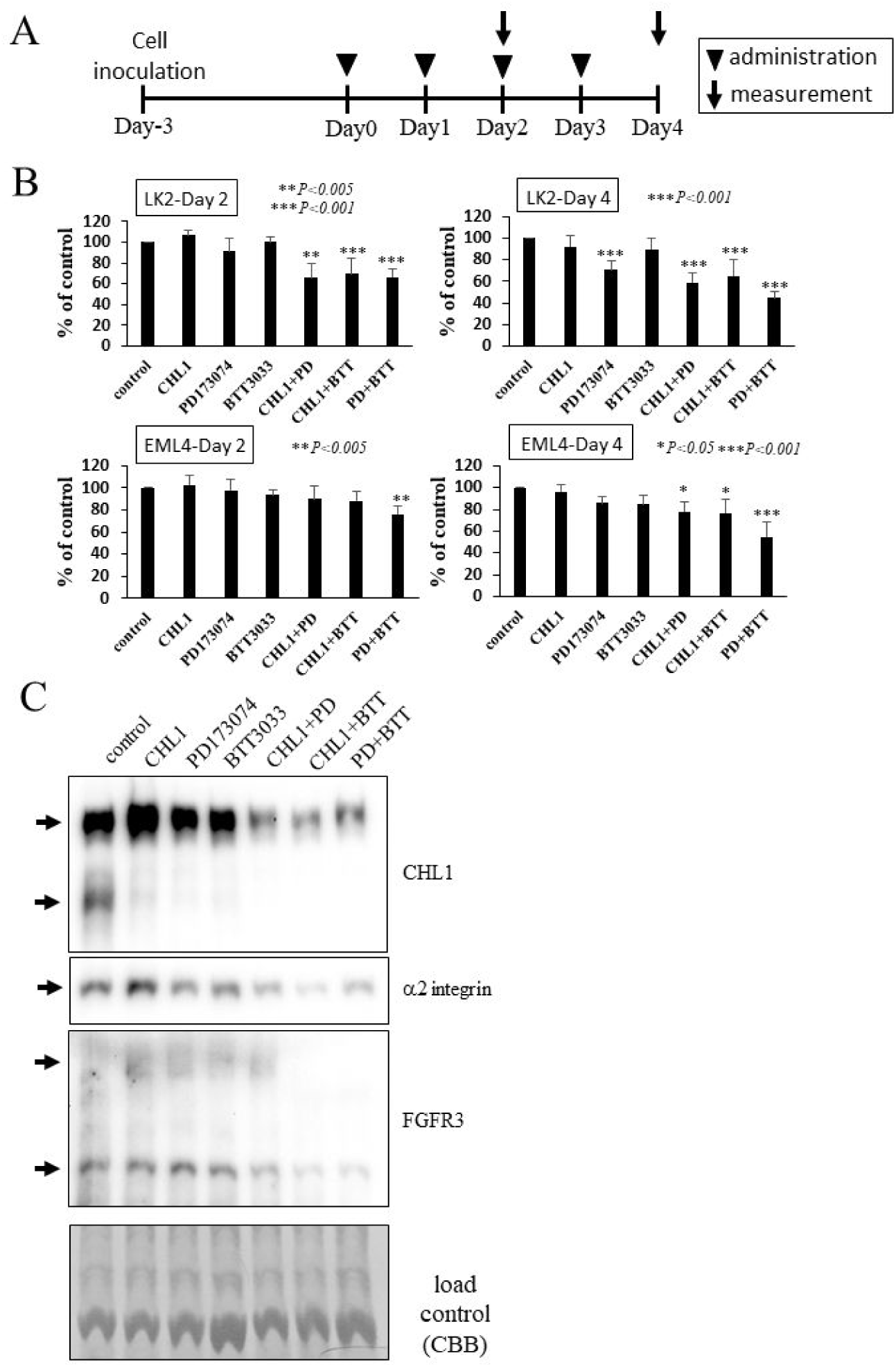
*In vitro* simulation of effective drug combination to inhibit cancer cell proliferation based on BiCAT information. (*A*) The single and double administration under daily treatment protocol (n = 5). The administration timing is indicated by closed triangles. The cell numbers of the treated cells were measured on Day 2 and Day 4. (*B*) The relative ratio (% of non-treated cells as control) of cell proliferation rates in LK2 cells and *EML4-ALK* primary cells. The statistical analysis was performed using Tukey’s and Dunnett's multiple test. The results from Dunnett’s test was in Fig. 6 \**P<0.05*; \*\**P<0.005*; *\*\*\*P<0.001*. (*C*) Double administration of molecular targeted reagents leads to changes in the expression and phosphorylation of partner molecules. The samples of single and double administration under daily treatment conditions (3 days) in LK2 cells were subjected to phos-tag SDS-PAGE and then western blot analysis using CHL1, α2 integrin, and FGFR3 antibodies. The CBB staining image indicates load control. The molecular weight markers were not shown in this figure because phos-tag SDS-PAGE cannot show the correct molecular weight of sample proteins.

As shown in Fig. S8A, double administration (CHL1+ PD173074, CHL1+BTT3033, or PD173074+BTT3033) was moderately effective (approximately 30% average inhibition) in contrast to single administration (approximately 10% average inhibition) for LK2 cells at Day 2 (Fig. S8A). Otherwise, double administration (PD173074+BTT3033) was only statistically significant for *EML4-ALK* primary cells at Day 2. The efficacy of double administration at Day 5 seemed to be greater variation or was weaker than that at Day 2 in both cell types. In daily treatments, as shown in Fig. 6B, double administration was similarly effective as single treatments, in contrast to no statistically significant inhibition after single administration for LK2 cells at Day 2. *EML4-ALK* primary cells at Day 2 showed similar results as cells treated with the single treatment. On Day 4, the double administration was clearly effective for LK2 cells (approximately 50% average inhibition) and moderately effective for *EML4-ALK* primary cells (approximately 30% average inhibition). In the every-other-day treatment condition (Fig. S8B), double administration was slightly less effective than the daily treatment protocol at Day 1 and Day 3 for LK2 cells except for PD173074+BTT3033 treatment. For *EML4-ALK* primary cells, double administration showed similar results as those with the daily treatment protocol at Day 1; however, there was significant efficacy in both single and double administration at Day 3.

These experiments under three protocols indicated that while the degree of inhibitory ratio differed among the protocols, we observed not only an additive effect but also a synergistic effect for the reagents against BiCAT molecules. For instance, the synergistic inhibitory effects of around 30% were observed at Day 2 by the double administration of three reagents in LK2 cells (Fig. 6B “*LK2-Day 2*”).

To examine the influence of double treatment, we performed western blot analysis on the reagent treated-cells. The UniProtKB database indicated that human and mouse CHL1, α2 integrin and FGFR3 are phosphorylated proteins (data not shown). Using Phos-tag^®^ gel, CHL1 was detected as two bands, which may reflect the degree of phosphorylation (Fig. 6D). The upper bands were reduced in only double administration samples (CHL1+ PD173074, CHL1+BTT3033 or PD173074+BTT3033) compared with the control and other single administration samples, in spite of equal amounts of loaded samples. In addition, the upper band in the sample with PD173074+BTT3033 double administration shifted upward compared with other upper bands, indicating that the phosphorylation of CHL1 was accelerated when double administration of PD173074 and BTT3033 was carried out in LK2 cells. As with CHL1, α2 integrin in the samples with double administration was reduced, whereas there was no significant shift of α2 integrin in the PD173074+BTT3033 double administration. FGFR3 was detected as two bands and was reduced in both CHL1+BTT3033 and PD173074+BTT3033 double administration samples without a significant band shift.

## Discussion

Various cancer treatments have recently been developed based on molecular targeted strategies (34). Here, we examined whether a BiCAT is useful as a novel cancer target for molecular targeted strategies.

The EMARS method that we previously developed (14) can be suitable for clarifying BiCATs in primary cancer cells under physiological conditions. We selected *EML4-ALK* transgenic mice for the study as the primary culture cells can be relatively easily established from the lung cancer tumor tissue derived from these mice. The establishment of primary culture cells from human cancer tissues has been reported (35). If primary culture cells can similarly be developed from human cancer biopsy tissue, BiCATs could possibly be simply identified using the EMARS method for each patient, resulting in personalized cancer medicine. Furthermore, the primary cancer cells are considered to be important for the *in vitro* simulation of medicine selection described in Fig. 6.

The selected target molecule for EMARS probe requires high expression and specificity in cancer tissue. It is, therefore, necessary to perform preliminary experiments (*e.g.* cDNA microarray) or pre-assessment for the decision of appropriate molecules. In the *EML4-ALK* transgenic mouse, CHL1 expression was restricted to the cancer tissue without any correlations to age and sex (Fig. 2B), suggesting that CHL1 seems to be good candidate target molecule.

In the proteomics analysis of EMARS products with mass spectrometry, there was no overlap of listed candidate molecules between *EML4-ALK* primary cells and LK2 cells (Table S2), indicating that the partner molecules constituting BiCATs with CHL1 were different among cancer cell types or species. However, we hypothesized that the partner molecule information obtained from *EML4-ALK* primary cells was applicable to LK2 cells and vice versa because it is sometimes insufficient to analyze due to differences in the ionization efficiency of molecules in mass spectrometry. Using cDNA microarray data, we also examined the changes of expression levels of representative partner molecules identified above. There was no significant change (data not shown), indicating that BiCAT formation was not simply dependent on the overexpression of both constituent molecules that occurs in cancer.

The immunostaining experiment of lung cancer tissues derived from lung cancer patients provides crucial information for the clinical application of BiCATs. Although it is a subjective analysis, approximately 40%–50% of cases were positive for three BiCATs (Fig. S7F). In lung cancer treatment, epithelial growth factor receptor (EGFR), including its mutant form, and *EML4-ALK* are well-known targets for molecular targeted lung cancer drugs such as Gefitinib (36, 37), Cetuximab (38) and Crizotinib (39). A previous study reported that EGFR is overexpressed in 40%–80% of non-small cell lung cancer patient (40), and EGFR mutations have also been detected in 19.4% lung cancer patients (41). The *EML4–ALK* fusion gene was detected in 6.7% of non-small cell lung cancer patients (25). The positive rate of these BiCATs in our study was comparable to typical cancer targets including tumor markers used for the index of lung cancer treatment. It is, therefore, possible that BiCATs contribute to diagnosis using pathological specimens from cancer patients, which has recently attracted attention as an effective molecular medicine strategy. Interestingly, due to the localization of BiCATs in specific cell populations of human lung cancer tissues (Fig. 5), BiCATs in pathological specimens may be used as a benchmark for a specific group of cells targeted for medicinal treatment, *e.g.* cancer stem cells (42), if we are able to identify the specific BiCATs expressed in these cells.

Regarding the cancer specificity of BiCAT, even if the BiCATs are determined by the EMARS method, appropriate BiCATs should be selected for further application. If there are 1,000 molecules present in a cancer cell membrane, there are theoretically approximately 500,000 BiCAT candidates because of the combination between two molecules. This provides a great opportunity for discovering new cancer targets, but it becomes difficult to select specific and appropriate BiCATs. Moreover, the existence of BiCATs in other normal tissues should also be assessed to determine specificity, which is crucial for effective molecular target strategies. The appropriate partner molecules constituting BiCATs should not be co-expressed in normal tissue. Using gene or protein expression databases, we can partially assess whether the candidate molecules are co-expressed in same normal tissues (Fig. S3 and S4). Whereas the database diversity prevented complete determination, some candidate combinations, for instance CHL1-α2 integrin, CHL1-FGFR3 (*EML4-ALK* primary cells), CHL1-α2 integrin, CHL1-FGFR3 and CHL1-contactin-1 (LK2 cells) were considered suitable BiCATs. In addition, it should be considered that partner molecules that form BiCATs with CHL1 are highly likely to form BiCATs with each other. In this study, we may also have to consider the combinations among FGFR3, α2 integrin, and contactin-1.

Moreover, it would be interesting if BiCAT information could contribute to the development of bispecific antibody medicine (43, 44) in recognizing cancer-specific BiCA for cancer treatment. In fact, strategies with antibodies recognizing two cell membrane molecules have been already developed as a molecular targeted bispecific antibody medicine for cancer treatment (43, 44). Among them, the bispecific antibodies recognizing *cis-*bimolecules similar to BiCAT have only been reported in a few cases (EGFR-IGFR (45), EGFR-Met (46), CD20-CD22 (47)). These anticancer targets are well known and predictable. In our strategy, it is an advantage that these can be further extended to the combination by typical membrane molecules. In addition, our strategy can identify multiple partner molecules and is also useful for Trifunctional antibody technology (Triomabs) (48), which has recently been drawing attention.

As a new approach to molecular targeted strategy, we attempted to perform a simulation of effective drug combinations for multiple drug administration (33) that inhibit cancer cell proliferation based on BiCAT information. This is based on previous findings that molecular complexes are important for signal transduction involved in cell functions through affecting other signals (49). For instance, CHL1 and integrins cooperatively contribute to signal transduction by interacting with each other (50–52). Although our results could not completely demonstrate whether bimolecular interactions in BiCAT contribute to the synergistic action of each reagent, BiCAT information has the possibility to help inform drug selection for multiple drug therapy with synergic effects. The efficacies of double administration in this study were not so powerful (especially for *EML4-ALK* primary cells), and thus it seems necessary to improve the selection of appropriate BiCATs by further studies. The molecular mechanism of this synergistic inhibition based on BiCAT information is unclear. Our results suggest that the molecules constituting the BiCAT are associated in proximal positions on the cell membrane so that the expression and phosphorylation of each molecule may be controlled via common upstream signal pathways, similar to that in lipid raft (49).

In conclusion, BiCATs have specific features and advantages in terms of the possibility of the development of novel targets and the improvement of antigen specificity not present in typical cancer targets, and may contribute to the discovery of effective and novel molecular targets.

## Experimental procedures

Part of the “*Experimental procedure*” are in the Supporting Information.

### cDNA microarray

mRNAs were purified from cancerous and non-cancerous parts of tissue from *EML4-ALK* transgenic mice using the RNeasy mini kit 74106 (QIAGEN). The purified mRNA samples were examined by an Agilent 2100 bioanalyzer (Agilent Technologies) to assess purity and concentration. Each mRNA sample was converted to cDNA with Cy3 (non-cancerous part) or Cy5 (cancerous part) labeling using the Agilent RNA Spike-In kit, two-color (Agilent Technologies) and Quick Amp labeling kit, two-color (Agilent Technologies), followed by purification with an RNeasy mini kit 74106 (QIAGEN). The labeled cDNA samples were mixed with hybridization solution in the Gene Expression Hybridization kit 5188-5242 (Agilent Technologies). The mixed samples were used in the Whole Mouse Genome Microarray Kit, 4×44K (G4122F: Agilent Technologies) and then incubated at 65°C for 17 h. The cDNA microarray was gently washed using the Gene Expression Wash Pack 5188-5327 (Agilent Technologies). The hybridized DNA microarray was scanned using Scanner G2505B (Agilent Technologies). The data was digitized and analyzed using Feature Extraction ver. 9.5.3.1 and GeneSpring GX ver. 11.5 (Agilent Technologies). Microarray expression data were deposited in Gene Expression Omnibus (NCBI) under the accession number GSE94261.

### Preparation of HRP-conjugated antibody for EMARS reaction

The human and mouse anti-CHL1 antibodies were partially reduced and bound to HRP using a peroxidase labeling kit SH (Dojindo). Anti-HRP antibody (Jackson Immunoresearch) was labeled with FITC (Sigma) and Alexa Fluor 647 (Invitrogen) for the validation as described below. The prepared HRP-conjugated CHL1 antibody was validated as follows: *EML4-ALK* primary cells and LK2 cells were incubated with HRP-conjugated antibody and then with the appropriate secondary antibodies, fluorescein-conjugated anti-HRP antibody (for LK2 cells) or Alexa Fluor 647-conjugated anti-HRP antibody (for *EML4-ALK* primary cells). Cells were observed with a confocal laser scan microscopy as described in Supporting experimental procedures.

### EMARS reaction for cell membrane

The EMARS reaction and detection of EMARS products were performed as described previously (14). Briefly, *EML4-ALK* primary cells and LK2 cells were washed once with PBS at room temperature and then treated with either 5 μ/ml of HRP-conjugated anti-mouse CHL1 and anti-human CHL1 antibodies or 4 μ/ml of HRP-conjugated CTxB (LIST Biological Laboratories) in PBS at room temperature for 20 min. The cells were then incubated with 0.1 mM fluorescein-conjugated arylazide or fluorescein-conjugated tyramide (15) with 0.0075% H_2_O_2_ in PBS at room temperature for 15 min in dark. The cell suspension was homogenized through a 26 G syringe needle to break the plasma membranes and samples were centrifuged at 20,000 g for 15 min to precipitate the plasma membrane fractions. After solubilization with NP-40 lysis buffer (20 mM Tris-HCl (pH 7.4), 150 mM NaCl, 5 mM EDTA, 1% NP-40, 1% glycerol), the samples were subjected to SDS-PAGE (10% gel, under non-reducing conditions). Gels were blotted to a PVDF membrane, which was then blocked with 5% skim milk solution. The membranes were then stained with goat anti-fluorescein antibody (0.2 μ/ml) followed by HRP-conjugated anti-goat IgG (1:3000) for FT detection.

### Staining of pathological specimens from lung cancer patients

This study used a lung cancer patient tissue array (No. OD-CT-RsLug04-003; Shanghai Outdo Biotech) that contains lung carcinoma tissues derived from 55 lung cancer patients (30 male and 25 female cases, mongoloid); among the total 55 cases, 53 cases have both tumor tissue and matched control normal tissue and two cases have tumor tissue only (53, 54). The specimens were deparaffinized with xylene and 70- –100% ethanol. Antigen retrieval was carried out using L.A.B solution (Polysciences Inc.) at room temperature for 10 min. The slides were then gently washed with PBS, treated with 5% BSA-PBS for 30 min and stained with anti-human CHL1 antibody (4 μ/ml) for 40 min followed by Alexa Fluor 546-conjugated anti-rat IgG (Thermo Fisher Scientific) for 40 min. After the CHL1 staining, the samples were subsequently stained with anti-α2 integrin antibody (4 μ/ml), anti-contactin1 antibody (4 μ/ml) or anti-FGFR3 antibody (0.8 µg/ml) for 40 min, followed by Alexa Fluor 488-conjugated anti-rabbit IgG (Thermo Fisher Scientific) for 40 min. The mounting media containing anti-fade reagent (DABCO; 1, 4-diazobizyclo (2, 2, 2) octane; Sigma-Aldrich) and DAPI (Nacalai Tesque) was incubated with specimens before observation. The samples were observed with an LSM 710 Laser Scanning Confocal Microscope (Carl Zeiss) mounted on an AxioImager Z2 equipped with a Diode laser unit (405 nm/30 mW), Argon laser unit (458, 488, 514 nm/25 mW), He-Ne laser unit (543 nm/1mW) and He-Ne laser unit (633 nm/5 mW). The objective lenses were EC-PLAN NEOFLUAR 5x/0.16 and APOCHROMAT 20×/0.8 (Carl Zeiss). Image acquisition and analysis was carried out with ZEN 2011 software (Carl Zeiss). Raw images including differential interference contrast image were captured under the identical settings in the case of same experiments and then exported to TIFF files. The distinction of positive or negative expression of BiCA was subjectively performed based on the detection of clear merged signals in ×5 visual fields between CHL1 and a bimolecular partner (α2 integrin, contactin1 and FGFR3) in each patient tumor tissue (Fig. S7D).

### In vitro proliferation inhibition assay

*EML4-ALK* primary cells and LK2 cells were grown on 96-well culture plates (in the case of *EML4-ALK* primary cells, the wells were coated with collagen I). After 72 h, antibody and/or chemical inhibitors against CHL1, FGFR3 and α2 integrin were added to medium as follows: anti-mouse CHL1 antibody (AF2147; R&D systems; final concentration 2.5 µg/ml), anti-human CHL1 antibody (MAB2126; R&D systems; final concentration 2.5 µg/ml), FGFR inhibitor (PD173074; Cayman Chemical; final concentration 30 nM) (55) and α2β1 integrin inhibitor (BTT3033; R&D systems; final concentration; 150 nM) (56, 57). The final concentration of each reagent was determined based on previous reports (22, 55, 57) and the data from a pilot study using *EML4-ALK* primary cells and LK2 cells (employed highest concentration in the data; Fig. S6). After treatment, short-term culture (3 to 5 days), additional treatment and cell counting were carried out according to three protocols: *protocol i*) single treatment and cell counting at Day 2 and Day 5 (Fig. S5A); *protocol ii*) daily treatment and cell counting at Day 2 and Day 4 (Fig. 6A); *protocol iii*) every-other-day treatment and cell counting at Day 1 and Day 3 with additional treatment at Day 2 (Fig. S5B). Cell counting was performed using the Cell Counting Kit-8 (Dojindo) with VarioSkan Flash microplate reader (Thermo Scientific) at 450 nm. Each protocol was carried out in multiple independent experiments (*protocol i*: n = 6, *protocol ii*: n = 5, *protocol iii*: n = 4 (in the case of *EML4-ALK* primary cells: n = 3)).

### Database search

The gene expression profile of mouse and human organs and tissues was obtained from BioGPS database (http://biogps.org/). The mouse and human datasets of BioGPS used in this study were GeneAtlas GNF1M, gcrma (mouse) and the GeneAtlas U133A, gcrma (human). The expression value of each gene was calculated from the average of the expression raw data derived from each probe sets (each raw data unit includes 1 to 4 samples). The protein expression profiles provided by proteome experiments were obtained from the Human Protein Atlas (http://www.proteinatlas.org/) and Human Proteome Map (http://www.humanproteomemap.org/).

### Data availability

The information on data availability in this study is summarized in Table S8.

### Statistical analysis

Statistical analyses were performed with Dunnett's multiple test (comparison to control cells) and Tukey's multiple test using R software (The R Foundation for Statistical Computing, Austria) and EZR (Saitama Medical Center, Jichi Medical University, Japan) (58), which is a graphical user interface for R. We used a statistical significance level of 0.05 or smaller. The statistical analyses indicated in Fig. 6 were performed by Dunnett's multiple test. The results of Tukey’s test are summarized in Table S7.

## Acknowledgements

We thank Professor Hiroyuki Mano and Dr. Manabu Soda for providing *EML4-ALK* transgenic mice; Professor Tsumoru Shintake for helping with electron microscopic analysis; and Kochi University experimental training equipment facility and Saitama Medical University Biomedical Research Center for technical assistance. The human squamous cell lung carcinoma cell lines LK2 (RCB1970) and RERF-LC-KJ (RCB1313) were provided by the RIKEN BRC through the National Bio-Resource Project of the MEXT, Japan. This work was supported by Grants-in-aid for Scientific Research in Japan (No. JP24590082, No. JP15K07941 and No. JP18K06663) (to N. K.), Mizutani foundation research grants (to N. K.), and Japan Science and Technology (JST) grants (to N. K.). We also thank Edanz Group (www.edanzediting.com/ac) for editing a draft of this manuscript.

## Conflict of Interest

The authors declare that they have no conflicts of interest with the contents of this article.

## Author Contributions

Conceptualization, N.K. and K.H.; Methodology, N.K., A.Y., and K.H.; Investigation, N.K., A.Y., T.O., R.K, Y.N., and Y.I.; Resources, R.K. and T.S., T.M., and

K.H.; Data Curation, N.K.; Writing-Original Draft, N.K., and K.H.; Writing - Review & Editing, R.K., T.S., T.N., T.M., and K.H.; Supervision, N.K., and K.H.; Funding Acquisition, N.K. and K.H.

